# High versatility to meet conservation targets for biodiversity and hydrological services at the Riviera Maya, Quintana Roo, Mexico

**DOI:** 10.1101/2023.03.23.533929

**Authors:** Juan Alberto Aguilar-Sánchez, Melanie Kolb

## Abstract

The ecosystem services (ES) framework has been proposed as an alternative to face the multiple challenges presented by biodiversity conservation, but the spatial conservation priorities of ES have been found to show low concordance levels with areas of high importance for biodiversity, which can lead to conflict during reserve design. To address this problem, the use of quantitative methods derived from systematic conservation planning has been proposed to identify spatial solutions that achieve the simultaneous representation of both elements in a spatially efficient manner. The aim of this study is to evaluate the differences between priority sites for biodiversity and hydrological ecosystem services (HES) using spatial prioritization models and to identify opportunities for co-benefits that allow an efficient conservation planning proposal, using as a case study the Riviera Maya, Mexico. The following hypothesis were tested by comparing models based on the prioritization algorithm Marxan: (1) Priority sites for biodiversity and HES are different, (2) HES priority sites adequately represent biodiversity conservation targets, and (3) integrating HES and biodiversity into one model is more efficient for representing conservation targets than combining the individual models for both elements. The results confirm: (1) Biodiversity and HES priority sites have different spatial patterns, sharing only 24% of priority sites, (2) HES priority sites achieve a high percentage (95%) of biodiversity conservation targets, showing that they can potentially be used for biodiversity representation, and (3) integrating HES and biodiversity into one model is more efficient to represent conservation targets than considering both elements individually (46% vs 66% of the study area). As there are no irreplaceable sites for biodiversity conservation, and less than 8% of the study area is covered by protected areas, there are clearly opportunities to align biodiversity and HES conservation actions at the Riviera Maya, Mexico. Despite the high context dependency of the spatial distribution of priority sites for biodiversity and HES, this study shows that the integration of conservation targets of both in the planning process can provide a solution to represent a high number of biodiversity and HES conservation targets.

## Introduction

A major potential for increasing biodiversity conservation is the possibility to consider ecosystem services (ES) in the conservation planning process, as both intrinsic and anthropogenic values of biodiversity and ES can be used to steer decisions towards biodiversity conservation with more diverse tools and actions [1-6]. But the combination of criteria for biodiversity and ES conservation will produce different outcomes, compared to only biodiversity, as human influenced or even transformed areas can be important for ES but not for biodiversity [7-9]. This not only means that priority areas for biodiversity and ecosystem services may be spatially distinct but also have varying levels of achieved conservation targets, efficiency, compatibility, and complementarity, depending on the context [1,10-16]. These differences are more evident at finer spatial scales, where the economic value and other benefits of ES are generally not concurrent with biodiversity [1], putting an emphasis on considering ES supply and demand areas in biodiversity conservation planning [7,13,17].

In the last decade, tools from the systematic conservation planning have been implemented with the aim of representing ES and biodiversity conservation targets in a spatially efficient solution, so that multiple objectives can be achieved during reserve design [1,18]. Systematic conservation planning optimizes conservation networks through an iterative identification of a smaller set of territorial units required to meet conservation targets, thereby reducing costs associated with conservation by selecting sites that are complementary to one another [19,20]. If a complementary set of sites can be identified, they could ensure the representation of biodiversity and ES, even if their respective priority areas have little spatial overlap [1,16,19].

Among the different types of ES, it has been observed that hydrological ES (HES) [21,22], in particular water supply and water flow regulation, are more spatially concordant with priority sites for biodiversity conservation [1]. Usually, HES are recognized as critical but not considered in decision making [6]. This tendency tends to be stronger regarding subterranean HES [23-27] because their degradation is not easily observed and thus are not considered a priority for conservation [23,26]. In Mexico, this is the case of the Yucatan Peninsula (YP), one of the largest groundwater reserves in the world, which is currently affected by the recent massive economic growth in a *karst* landscape with low natural filtration capacity [28]. The high hydraulic conductivity of the YP’s limestone platform makes the aquifer not only highly susceptible to contamination but also the only source of drinking water in the region [28-31]. In particular, in the Riviera Maya, a major tourist corridor on the east coast of Quintana Roo, population growth and urban development have notably increased the degradation of water resources [30,32].

Due to the economic importance of HES and biodiversity in the YP, conservation planning for groundwater protection must consider the ecohydrological processes that secure the regional water supply and biodiversity maintenance [28,33-35]. Finding spatially efficient solutions in reserve design intended to represent both elements would be important to facilitate conservation actions in the Riviera Maya. Besides, the consideration of HES opens the possibility to integrate terrestrial and aquatic biodiversity conservation targets as they consider ecohydrological aspects, like the aquatic-terrestrial interface in conservation planning [22,27,35,36].

The aim of this study is to contribute to the understanding of the effects of including ES in the process of establishing conservation priority sites by evaluating the differences between priority sites for biodiversity and ES using spatial prioritization models and identify opportunities for co-benefits that allow an efficient conservation planning proposal. The following hypothesis were tested: (1) Priority sites for biodiversity and HES are spatially different, (2) HES priority sites adequately represent biodiversity conservation targets, and (3) integrating HES and biodiversity into one model is less efficient to represent conservation targets than considering both elements individually.

## Materials and methods

### Study area

The case study to test these hypotheses is the Riviera Maya, Quintana Roo, Mexico, which forms part of the YP, a 165,000 km^2^ limestone platform located in southeastern Mexico [28] (Fig 1). The dissolution of the limestone allows the formation of interconnected tunnels and conduit systems of different sizes including the world’s largest underwater cave system [28,37-40]. The *cenotes*, as sinkholes resulting from cave collapses are locally called, together with the high permeability of the limestone represent a conduit for above ground materials and energy, sustaining unique underground ecosystems, many of them anchialine, with high diversity of endemic *stygobiont* species [29,31,41-44].

**Fig 1.**
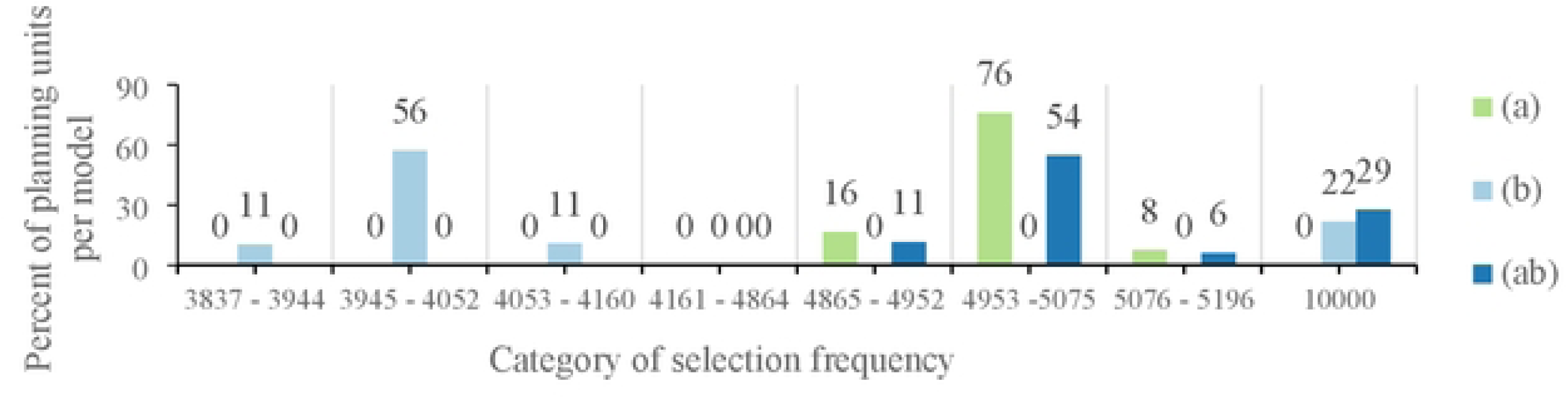
Location of the study area. Localities and the planning units used in the prioritization analysis are shown.

Epigean and hypogean ecosystems depend on the interaction between the water and limestone [35,44]. The *karst* system maintains, through the hydrological connections, many processes that support different ecosystems from tropical forests to wetlands and coral reefs, creating a functional transversal corridor from terrestrial ecosystems to coastal lagoons and estuaries [34]. Depending on the scale of these connections, different processes and ecosystem functions are maintained [28,35,40,43].

The beauty of the *karstic* features and the Caribbean Sea, together with its varied ecosystems and cultural heritage, makes the YP one of the most important tourist centers in Mexico. Most of the current economic activity is centered along the east cost of Quintana Roo. This corridor is known as “The Riviera Maya” and includes the municipalities of Solidaridad to the north and Tulum to the south, and extends approximately 40 km inland to the border with the Yucatan state [28,32].

The exponential demographic and economic growth of the last three decades has had a negative impact on the environment due to the land use change and the intrinsic geological vulnerability of the aquifer to contamination [28,31-33,45-47]. Water quality in the YP has been decreasing with important health risks for humans and ecosystems, particularly at the Riviera Maya where currently sewage treatment is non-existent [31,47-51]. Together, all these factors pose an urgency for advancing conservation planning [28,32].

Given the difficulty to delimitate watersheds in the YT due to a lack of topographic features, the study area corresponds to the central zone of the *Holbox* fracture system [52] with an area of 10184.6 km^2^ (Fig 1). The main vegetation types are medium evergreen forest and a variety of aquatic vegetation [53]. The most representative soil types are regosols and leptosols [54].

### Planning units

Three sizes of planning units (PU) were established according to the levels of landscape heterogeneity (land cover types, settlements, roads) and distance from the coast using SDMtoolbox v2.5 for ArcMap 10.0 [55]. The smaller PU (500 m^2^) covered the first 5 km from the coast, the medium PU (1 km^2^) inland in areas with settlements and roads, and the larger PU (5 km^2^) in areas with higher land cover homogeneity (Fig 1). This spatial distribution of the 6595 PU (S1 Map) was chosen so that the different levels of biophysical and pressure factors in the different zones of the study area can be considered in the prioritization process.

### Conservation targets

#### Biodiversity

A total of 319 different spatial elements were used to represent biodiversity. They corresponded to 273 potential species distribution models (S2 Dataset), 38 georeferenced points of species occurrences (S3 Dataset), and eight surrogates. The species data was obtained from the CONABIO-SNIB portal [56], and the GBIF database [57]. Species of seven biological groups (crustaceans, fish, amphibians, reptiles, mammals, birds, and plants) were selected based on different attributes of their distribution and conservation status [58-61]. The surrogates corresponded to physiographic elements that potentially can represent biodiversity at a regional level [62], in this case by the natural vegetation types [63], water bodies [63], and a 15 km range from the coastline inland, which represents critical habitat for subterranean fauna [64] (S4 Table).

Quantitative conservation targets express the “conservation goals” to be achieved for each conservation element [19,62]. In this study, they correspond to the percentage of the total area of each biodiversity or ES spatial elements that should ideally be represented within the priority sites. For species data, conservation targets were based on endemicity, rarity (species that occupied the last quartile of the geographic distribution range of each taxonomic group), the extinction risk status according to the Mexican red list (NOM059 [58]) and the international red list (IUCN [59]), the pressure from international commerce (CITES [60]), and species with national priority degree (CONAMP [61]) (Table 1).

**Table 1.** Examples of conservation goal allocation according to biodiversity criteria values. [Abbreviations: YP= Yucatan peninsula; Mex= Mexico; IUCN= International Union for Conservation of Nature; E= possibly extinct in the wild; P= at risk of extinction; A= threatened; Pr= subject to special protection; Cr= critically endangered; En= endangered; Vu= vulnerable; Lc= least concern; CITES= Convention on International Trade in Endangered Species of Wild Fauna and Flora; Ap= appendix; VH= very high; H= high; M= regular; Biodiversity criteria values are shown on the table heading]. *The rarity was evaluated using as a threshold the last quartile of the geographic distribution range of each taxonomic group at the national scale models. This criterion does not apply to the species represented by point occurrences because the distribution was assumed as the total area of the PU in which they occurred.

|  | **Endemicity**<br>**YP/Mex**<br>**25/12** | **Mexican**<br>**red list**<br>**(NOM-059)**<br>**E/P/A/Pr**<br>**20/20/10/5** | **IUCN**<br>**red list**<br>**Cr/En/Vu**<br>**10/7/5** | **CITES**<br>**ApI/ApII**<br>**10/5** | **National**<br>**Priority**<br>**VH/H/M**<br>**10/10/5** | **Rarity\***<br>**1st**<br>**/other**<br>**10 / 0** | **Score** | **Assigned**<br>**target** |
| --- | --- | --- | --- | --- | --- | --- | --- | --- |
| **Species 1**<br>**(distribution model)** | 25 | 20 | 10 | 10 | 10 | 10 | 85 | 50% |
| **Species 2**<br>**(point occurrences)** | 12 | 5 | 5 | 5 | 5 | NA | 32 | 20% |

\*The rarity was evaluated using as a threshold the last quartile of the geographic distribution range of each taxonomic group of the national distribution. This criterion does not apply to the species represented by georeferenced points because the distribution was assumed as the total area of the PU in which they occurred.

For distribution models, conservation targets were based on the sum of values assigned to each conservation criteria as follows: 10% (Σ = 1–21); 20% (Σ = 22 - 41); 40% (Σ = 42 - 63); 50% (Σ = 64–85) (Table 1). In the case of point occurrences, all the PU with a data point were considered as the distribution area and conservation targets were assigned as follows: 10% (Σ = 1-18); 20% (Σ = 19-37); 40% (Σ = 38 - 56); 50% (Σ = 56-75) (Table 1).

In the case of vegetation types, conservation targets were established based on their relative distribution area: 50% (<2% of the study area); 30% (>=2 and <5% of the study area); 10% (>5% of the study area). For water bodies, an intermediate value of 30% was established. For the critical habitat zone for subterranean fauna, a value of 40% was established, due to the extremely high human pressure.

#### Hydrological ecosystem services

HES were defined as the benefits produced by the effect of ecosystems on groundwater, referring to the ecohydrological processes and biophysical attributes that secure the regional water supply in terms of provision (amount of water) and regulation (water quality) [21,22,27]. HES demand was represented by the water concessions in the Public Registry for Water Rights (REPDA [65]). For HES supply, 12 spatial elements related to the areas that contribute a greater amount of water to the aquifer through infiltration, were included in this study: Continental water bodies [63], regional-scale fractures [66], the *Ox Bel Ha* and *Sac Actun* cave systems [39], an 8 km range from the coastline inland that represent the coastal areas with an extremely high aquifer geological vulnerability [67], areas with the highest infiltration capacity based on a water balance (precipitation from [68], minus the actual evapotranspiration from [69]; in all cases, only the highest quartile was considered as a conservation element (S5 Map). For fractures, underwater cave systems, and vulnerable aquifer areas, an intermediate value of 20% was established. For the area with the greatest infiltration, a conservation target of 30% was established.

The conservation targets for land cover types were established considering 1) total infiltration (l/km^2^), 2) water concessions (l/km^2^), and 3) the total extent of land cover type (km^2^). All resulting values were ranked for each criterion (in the case of the total surface from major to minor values) to establish quartiles to which to assign a conservation target: First quartile = 2.5%, second quartile = 5%, third quartile = 7.5%, fourth quartile = 10%. The final conservation targets were calculated as the sum of the % of all criteria (S6 Dataset).

Considering the extremely high hydrologic connectivity and the importance of ecohydrological interaction between the different types of vegetation and the hydrological system, a conservation target was assigned to all natural land covers [63], representing the average of the conservation targets for each vegetation type.

### Prioritization analysis

#### Models and parameters

Three models were generated using Marxan v.2.4.3 [70], one for biodiversity elements (a), one for HES (b) and one considering both (ab). Marxan is designed to resolve the minimum set problem to select PU in an iterative process that allows prioritizing spatial solutions with PU that better complement each other representing a higher number of conservation targets. For each model, 10000 runs with 1000000 iterations were performed. A threshold of minimum 90% of targets, a proportion of 0.4 of the initial sampling and a border length of 1 were specified. The selection frequency allows evaluating whether a site can be replaced by another to meet the conservation targets of a region and is an indicative value of the relative importance or irreplaceability of that site, so it can be used to delimit priority sites [71,72].

In model (ab), shared conservation elements were revaluated: An average of the HES and biodiversity conservation targets was assigned (in the case that the biodiversity conservation targets were higher) or half of the biodiversity target was added to the HES target (in the case that HES conservation targets were higher) (S6 Dataset).

#### Pressure factors and demand areas

For the models (a) and (ab) information on human pressure factors on biodiversity was used to establish conservation priorities based on a proactive approach [73]. A map of pressures factors was created including the following criteria: Urban areas [63], the inverse ecological integrity [74], human population density [75], and agricultural areas [63], road density [76]. Each factor was weighted, and the corresponding values were summed for each PU (S7 Table, S8 Dataset).

For (b) and (ab) models, the PU with the presence of water concessions were considered as HES demand areas and fixed *a priori* in the initial selection, as they serve to spatially connect HES supply and demand [7] (S9 Dataset).

#### Site hierarchy and null models

To obtain priority levels, the PU from the best solution identified by Marxan were ranked and classified based on their selection frequency for each model. Then the accumulative area and the accomplished conservation targets were plotted and where the maximum number of conservation targets was achieved, this subset was divided into quartiles. PUs in the first quartile were considered as extreme priority sites, the second as highly important, the third as medium priority sites for conservation. The rest of the study area was considered as low importance.

The spatial efficiency of the models was evaluated by a null model, taking as a reference the area covered by the obtained prioritization models (37% for biodiversity and 45% for HES), but with randomly selected planning units. This way, the null model represented the average conservation targets achieved running 10000 random solutions (S10 Dataset for biodiversity and S11 Dataset for HES).

#### Gap analysis

Finally, the resulting priority sites were evaluated regarding their representation in existing protected areas [77].

## Results

### Model features and achieved conservation targets

The models (a), (b) and (ab) cover 37, 45% and 46% of the study area respectively (Table 2). The union of models (a) and (b), is equivalent to 66% of the study area as they only share 24% of shared priority sites. Despite the low spatial congruence between biodiversity and HES conservation priorities, the results also show that most PU obtained low or intermediate selection frequency values in all models (<5,000), which means that there is ample opportunity for the prioritization algorithm to accomplish a very high number of conservation targets (Fig 2). The identified irreplaceable sites (selection frequency = 10,000), represent 9% of the study area and corresponded to the demand areas, which were established *a priori* as a priority sites in models (b) and (ab).

**Fig 2.**
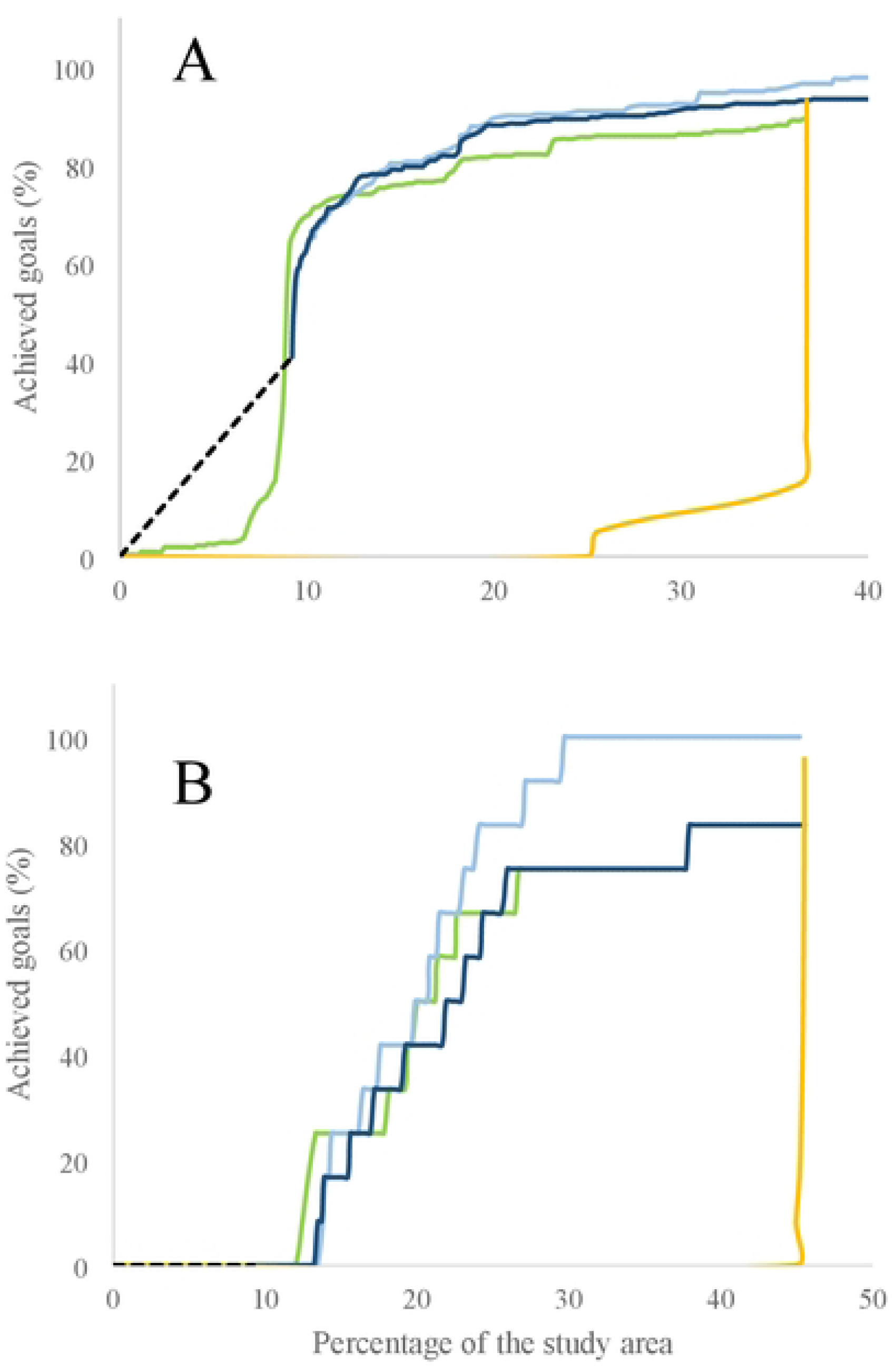
Selection frequency distribution by model. This data is derived from planning units of all solutions found by Marxan.

**Table 2.**
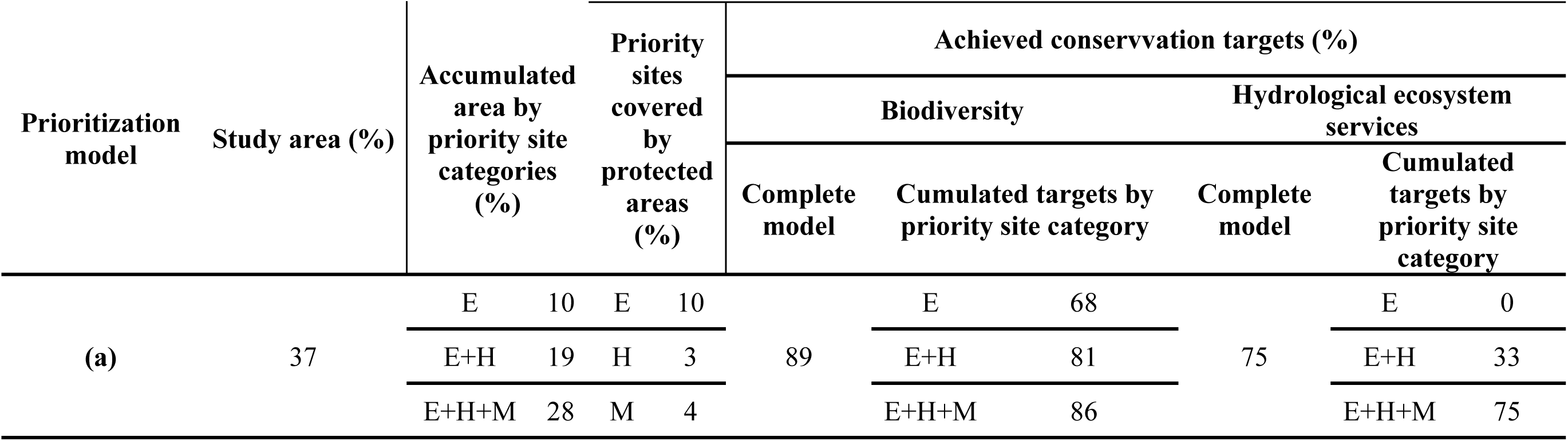
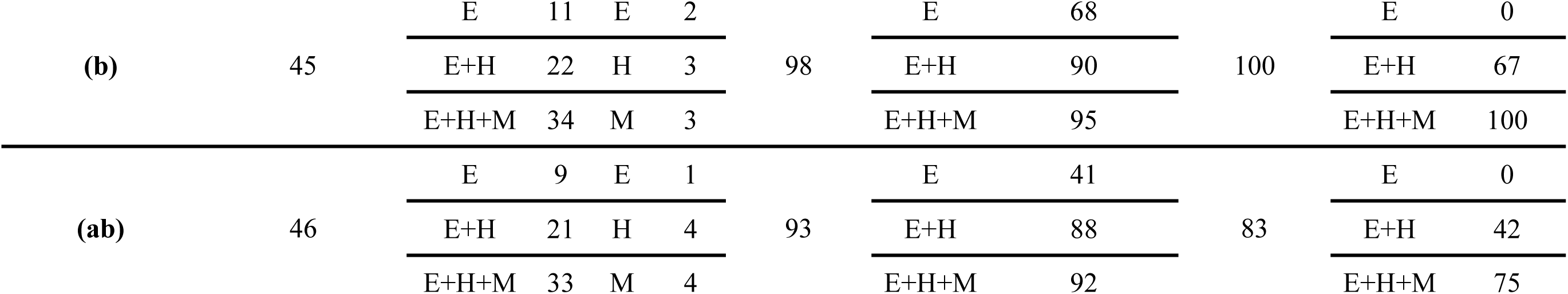
Marxan models features and priority site representation in regional protected areas. [Abbreviations: E= Extreme priority sites, H=High priority sites, M= Medium priority sites].

Model (b) achieved the highest percentage of conservation targets for both, biodiversity (95%) and HES (100%), while model (a) achieved the lowest representation values (75% and 86% for HES and biodiversity, respectively) (Table 2). Model (ab) showed an intermediate level of achieved conservation targets for both types of criteria (Table 2). The accumulation curves show that all prioritization models have similar trends (Fig 3). Model (a) accumulates biodiversity conservation targets faster than models (b) and (ab) before it reaches 12% of the study area (Fig 3). The model (ab) accumulates a higher number of biodiversity conservation targets between 12% and 14% of the study area and the model (b) achieve more conservation targets than the others models once it exceeds that range. The accumulation of accomplished biodiversity conservation targets was quicker than for HES in all models. But HES demand areas are comparatively less efficient in accumulating biodiversity conservation targets than model (a) (41% vs 64% in 9% of the study area respectively) (Fig 3). Most (>50%) biodiversity conservation targets can be achieved using 9% of the study area, while a minimum of 21% is needed to adequately represent HES conservation targets (Table 2). Extreme, high and medium priority sites need to be considered to achieve most HES conservation targets, with the exception of model (b): extremely and highly important sites achieve 66% of the HES conservation targets (Table 2). Extreme priority sites of the models (a) and (b) are similar in size (10% and 11% of the study area respectively) and achieve the same number of biodiversity conservation targets (66% of objectives), showing that both can potentially be used for biodiversity representation (Table 2).

**Figure 3.**
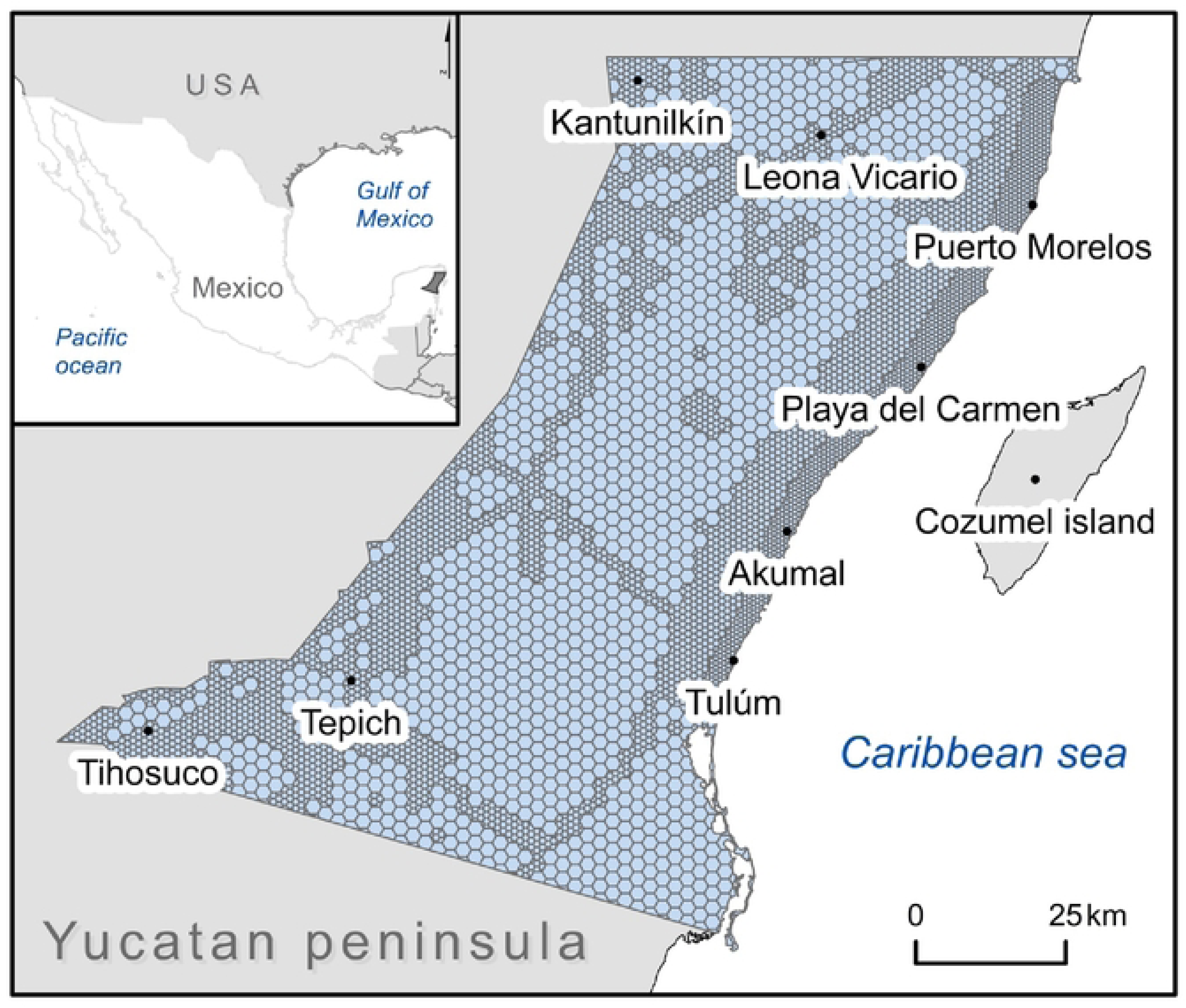
Effectiveness in meeting the biodiversity (A) and hydrological ecosystem services (B) conservation targets of the models as the study area is added (%). Accumulation curves derived from the best solution: green line: (a), light blue: model (b), dark blue: model (ab), yellow line: null model. The pointed line refers to the effect of demand areas set *a priori* as priority sites in models (b) and (ab).

The null models need much more area to achieve the same level of conservation targets as the other models, so its accumulation curves remain lower until reaching 36% and 45% of the study area for biodiversity and HES conservation targets respectively (Fig 3).

### Spatial pattern of priority sites

The priority sites for HES (*b*) present a more uniform distribution throughout the study area than in (a) and tend to cluster near the areas of demand (Figs 4a and 4b). The biodiversity priority sites tend to cluster near the central, N, and S parts of the study area and are located far from the coastal zone and the SW part, which is adjacent to the Yucatán state (Fig 4a). Thus, the spatial configurations of (a) and (b) show the greatest contrast in the coastal zone and in the NW, NE, and SW parts of the study area (Fig 4). Model (ab) shows intermediate spatial patterns between (a) and (b) models: While (ab) priority sites are uniformly distributed throughout the study area and conglomerate near the central and N parts, like what occurs in (b), they also are a less frequent in the SW part, more similar to model (a) (Fig 4c).

**Fig 4.**
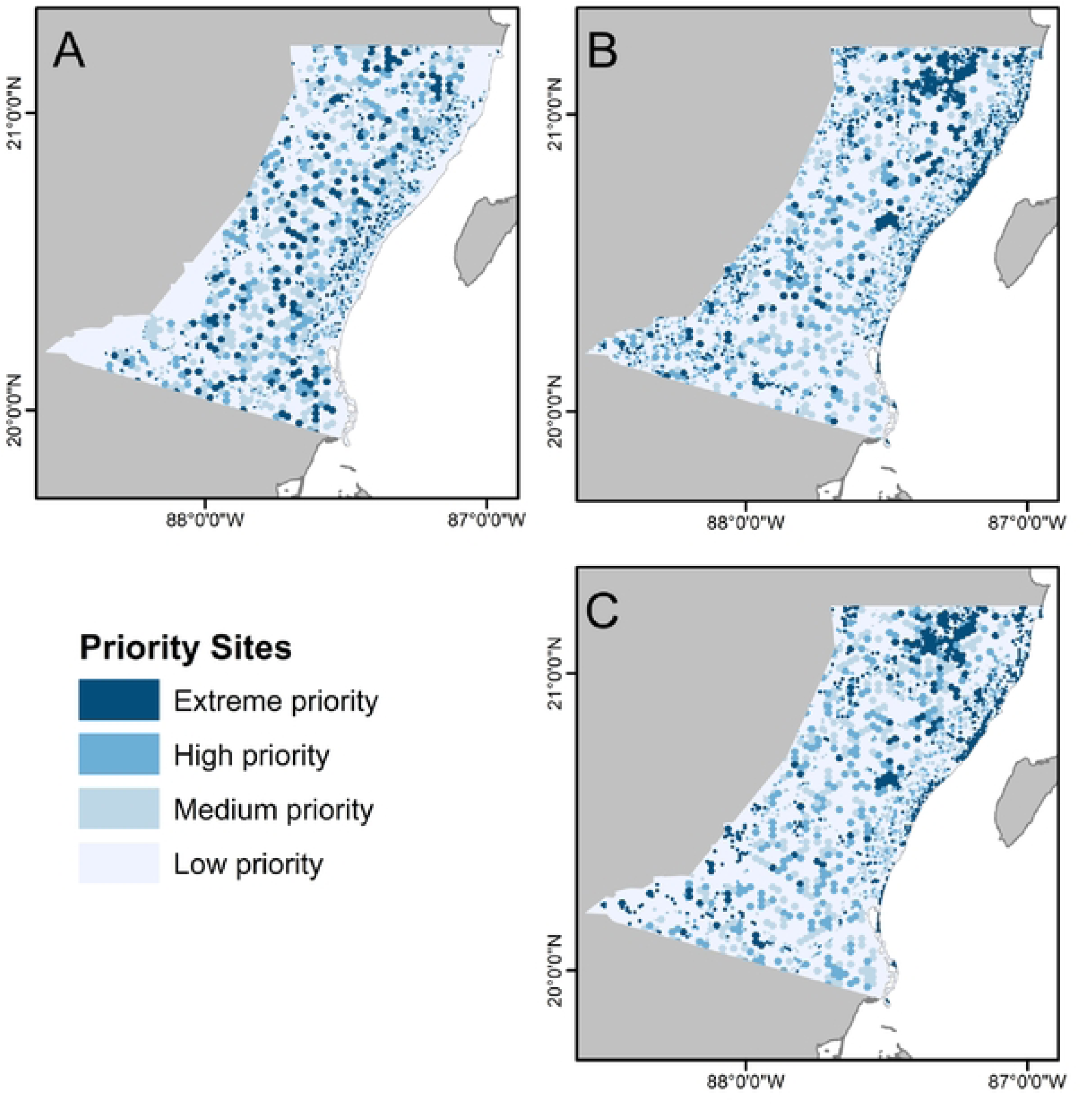
Priority areas for A) Biodiversity (a), B) Hydrological ecosystem services (b) and C) Combination (ab) based on the best solution proposed by MARXAN.

## Discussion

Even though the co-benefits between biodiversity and ES conservation have been proposed as the major objective of conservation [1,2,4,5,18,78], it is not clear how to organize this in a certain context [7,8]. Until now, the combination of both conservation priorities has achieved little attention, as spatial congruence tends to be low. In this study, while the results indeed confirm that there is a low spatial correspondence between conservation priorities, when seen separately, they also show that the combination of both criteria can be used to represent a high number of biodiversity and HES conservation targets. Thus, even if the combined priority sites are slightly less spatially efficient than priorities for each (46% vs 37% for biodiversity and 45% for HES), the main critique of using ES for biodiversity conservation, priorities for ES not being congruent with the ones for biodiversity [8] can be answered by an integrative approach, at least in certain contexts.

In general, the models with conservation targets for HES require more area and the conservation goals are achieved slower. But the fact that model (b) can potentially achieve more biodiversity conservation goals than model (a), clearly shows that it is possible to represent biodiversity using HES priority sites. The combination model is much more efficient than joining the individual models for biodiversity and HES (a ∪ b, 46% vs 66% of study area). Together with the low efficiency to achieve the conservation targets with randomly selected sites, these findings highlight the importance of combining biodiversity and HES criteria to meet multiple conservation objectives at the Riviera Maya.

Similar to the findings of Cimon-Morin et al. [1], the low spatial congruence between biodiversity and HES priority sites in this case study could be due to: 1) the little spatial correspondence between to the chosen conservation targets to represent biodiversity and ES (only 2% of conservation targets are shared in model (a) and (b)); 2) the use of HES demand areas as *a priori* set priorities in the model (b); and 3) the use of the human pressure factors map in model (a). Using demand areas *a priori* implies more cohesive and larger priority sites near human settlements, while the use of the human pressure factor map leads to priority sites far from the areas of greatest anthropic influence. Thus biodiversity representation in HES priority sites could be overestimated, as pressure factors tend to concentrate in HES demand areas, and the conservation of several groups like large, specialized, or sensitive species to human disturbance could be unavailable or not compatible.

While most biodiversity conservation targets (64%) can easily be represented in 9% of the study area, achieving the same level of representation of HES conservation targets would ideally require between 21 and 24% of the study area. This fact could make HES conservation more difficult to manage by requiring more resources. But at the same time offers the possibility to connect ES supply and demand to assure that ES can be realized, and benefits can be obtained. HES conservation planning ideally requires the spatial coupling of the areas where the ecological processes occur with the areas where the users who benefit from them are located [6]. In the case of model (ab), the HES flow areas that connect areas of supply and demand can be used to accomplish biodiversity goals, making HES priority sites a kind of umbrella to facilitate biodiversity conservation.

There is currently a general lack of knowledge about how biodiversity and hydrological processes interact, which can give the impression that there is little or no relationship between both components. In future evaluations, the spatial congruence between biodiversity and HES could be enhanced by including more biodiversity elements from aquatic ecosystems, but also regarding biotic regulators of the *karst* processes itself [35,38], as well as elements related to the maintenance of habitats, including in the marine realm [34,79-82]. There is still a lack of knowledge about the groundwater ecology and subterranean biodiversity in the region, despite the regional and national importance of the study area [41] and it is necessary to increase the knowledge on aquatic biodiversity, and especially *stygobionta*, to overcome the current limits for the inclusion of cross-system conservation targets.

Future studies also should consider that the spatial relationships between ES and biodiversity are location-specific, so that for multi-scale planning and decision-making needs to be implemented to reflect local and regional priorities [1,13,14]. In this context, to capitalize on co-benefits between biodiversity and HES in the Riviera Maya, finer scale analyses are needed for the coast, as it is particularly vulnerable to the degradation of hydrological resources due to the low natural filtration capacity of the highly porous limestone, the amount of resources demanded and waste produced by urban and touristic areas, and the saline intrusion that occurs due to the disturbance of the aquifer [40].

Given that there is a great versatility to meet biodiversity conservation targets in all models, HES-based conservation arguments in the Riviera Maya could be very useful to justify and facilitate financing conservation action [1-3,6,13]. In certain areas with a high level of conflict among biodiversity conservation and land use, like the Riviera Maya, the consideration of ES in conservation planning can mean a way forward to reconcile both objectives. For the YP, an increased rank for ES conservation, in regard to biodiversity priority has been found and in this context, a multipurpose approach could be implemented based on priority sites as an alternative to also promote and facilitate biodiversity conservation [2,3,5]. Biodiversity and ES loss need urgent attention from decision makers to assure their conservation and derived benefits for humanity. All models presented in this study provide a starting point to establish discussion about conservation priorities at the Riviera Maya, Mexico.

## Acknowledgments

We want to thank Dr. Patricia A. Beddows for her valuable feedback and guidance with the hydrology and geology part of this study. We also want to thank Dr. Fernando Álvarez from the National Crustacean Collection, who guided us in the conservation of anquihaline fauna.

## Supporting Information

**S1 Map. Map of planning units.**

**S2 Dataset. Conservation targets based on species distribution models. S3 Dataset. Conservation targets based on occurrence registers.**

**S4 Table. Biodiversity surrogates. S5 Map. Water balance model.**

**S6 Dataset. Hydrological ecosystem services conservation targets. S7 Table. Pressure factors on biodiversity criteria.**

**S8 Dataset. Weighted pressure factors values by planning unit. S9 Dataset. List of the planning units and their features.**

**S10 Dataset. Null model of achieved conservation targets for biodiversity.**

**S11 Dataset. Null model of achieved conservation targets for hydrological ecosystem services.**

